# IL-6 trans-signaling regulates CXCL12 expression in mesenchymal stromal cells and plasma cell niche in rejecting lung allografts

**DOI:** 10.1101/2025.02.16.638532

**Authors:** A. Patrick McLinden, Kazuki Tadokoro, Yuta Ibuki, Natalie M. Walker, Lu Lu, Fang Ke, Ragini Vittal, Aditi Ranjan, Monica P Mehta, Carol F. Farver, Joshua D. Welch, Vibha N. Lama

## Abstract

A transplanted lung offers a permissive *milieu* for local adaptive immune cell responses. Here, we characterize a transcriptionally and anatomically distinct adult lung-resident mesenchymal stromal population (MSC) that supports a pro-survival niche for antibody-secreting cells in rejecting lung allografts through a novel IL-6 trans-signaling/CXCL12 axis. By using a mouse orthotopic lung transplant model and *Blimp1^EYFP^* recipients, we identify spatial localization of antibody-secreting cells (ASCs) and terminally differentiated plasma cells (PCs) along the bronchovascular bundles (BVBs). A previously described Foxf1^+^Gli1^+^Itga8^−^ subset of collagen-expressing MSCs, which forms a 3-dimensional network along the bronchovascular bundles (BVB-MSCs), was found to be the major source of the PC survival factors CXCL12 and IL-6. *Cxcl12^iCre/ERT2^Rosa26^tdTomato^*mice utilized as donors, validated the expansion of this population in a rejecting graft and their intimate association with ASCs. CXCL12 expression was increased in murine allografts and in Foxf1^+^ mesenchymal cells (MCs) isolated from human CLAD patients. IL-6 trans-signaling/STAT3 signaling axis was shown to upregulate CXCL12 secretion in human MCs, and Olamkicept-mediated neutralization of IL-6 trans-signaling in murine RAS attenuates CXCL12 expression, intra-graft ASC population, and fibrogenesis. Our findings represent the first delineation of specialized CXCL12-expressing mesenchymal stromal cells in adult lungs and the contribution of IL-6 trans-signaling driven CXCL12 expression to sustaining intra-graft ASC niches and allograft fibrogenesis.

**One Sentence Summary:** We characterize CXCL12-expressing mesenchymal cells and their role in a pro-survival niche for antibody-secreting cells in rejecting lung allografts.

## INTRODUCTION

Lung transplantation remains the only viable option for patients with end-stage lung diseases (*1*). However, it carries unique challenges, including rapid organ allocation that precludes HLA matching and an inherently high immunogenic environment with direct environmental exposure (*2, 3*). Long term survival after lung transplantation remains lowest among all solid organ transplants with chronic lung allograft dysfunction (CLAD) being the primary cause of mortality after one year (*4*). Failing lung allografts, like other chronically rejecting solid organs, exhibit immune cell infiltration and fibrosis, with a recalcitrant course marked by progressive decline (*4–6*).

Restrictive allograft syndrome (RAS) is a particularly fulminant form of CLAD, distinguished from bronchiolitis obliterans syndrome (BOS), the other common phenotype of CLAD, by its restrictive physiology on pulmonary function testing and characteristic histopathological and radiographic features (*7–9*). The development of RAS has been associated with donor-specific antibodies and antibody-mediated rejection in human studies, suggesting a role for humoral immune activation in its pathogenesis (*10–13*). We have recently characterized a novel orthotopic murine single lung transplant model of RAS that mimics human pathology (*14*). A comparison of RAS and BOS murine models demonstrated humoral immune skewing as unique to RAS (*14*). A requisite role for humoral immune cell activation in RAS pathogenesis was established with allografts transplanted into mature B cell-deficient (*µMt^-/-^*) or antibody-deficient (*AID^-/-^ s^-/-^*) mice demonstrating protection (*14*). Accumulation of activated B cells, plasmablasts, and fully differentiated plasma cells was a prominent and unique feature noted in the murine RAS lung allografts. Intra-graft PC infiltration and donor-specific antibodies (DSAs) have also been noted in human RAS lungs (*14, 15*). However, the temporal and spatial establishment of local antibody secreting cell (ASC) niches and their regulation in a rejecting allograft remains to be established.

MCs play a key role in chronically rejecting allografts, with fibrosis being a pathognomonic feature of CLAD (*16–19*). While airway-restricted fibrosis is noted in BOS, more robust fibrosis along bronchovascular bundles and in the subpleural areas is noted in RAS. Our prior clinical studies have brought attention to resident mesenchymal progenitors in human adult lungs by demonstrating donor origin of multi-potent mesenchymal cells isolated from BAL of gender mis-matched human lung allografts (*16*). Subsequently, we identified them as lung-specific cells with unique expression of embryonic lung mesenchyme-associated transcription factor Foxf1, and as key contributors to allograft fibrosis (*19*). Significant insight into allograft fibrogenesis came from studying MCs derived directly from human normal and CLAD lung allografts (*17, 20–25*). In recent work, we characterized the *in vivo* niche of Foxf1^+^ MCs and found that Foxf1 expression is retained in a transcriptionally distinct MC population that resides in a three-dimensional network along the bronchovascular bundles; Foxf1 was noted to be conspicuously absent in MCs in the alveolar compartment (*26*). Lineage tracing studies in the murine RAS model confirmed the expansion of these Foxf1^+^Gli1^+^Itga8^−^Col1a1^+^ cells in rejecting lung allografts (*26*). While our investigations have shed light on the fibrotic functions of these cells in CLAD, MCs are also known to play a key role in regulating the microenvironment through their paracrine actions, and are well-recognized as specialized stromal cells that colocalize and support local immune cell niches (*27–29*).

Given that rejecting RAS lung allografts provide a pro-survival niche for antibody-secreting cells (ASCs), we investigated the spatiotemporal establishment of these niches and the MCs contributing to it. We demonstrate that Gli1^+^Foxf1^+^Itga8^−^Col1a1^+^ bronchovascular bundle-associated MCs (BVB-MCs) are a primary structural component of the ASC niche and source of the key plasma cell recruitment and survival factors CXCL12 and IL-6. A novel mechanism of upregulation of CXCL12 expression in BVB- MCs by IL-6 trans-signaling and downstream JAK/STAT activation is identified. Of key significance is the demonstration that targeting IL-6 trans-signaling with Olamkicept inhibits ASC accumulation and mitigates allograft fibrosis.

## RESULTS

### Temporal establishment of pro-survival ASC niches in murine RAS allografts

We have previously shown that RAS phenotype of chronic allograft rejection is associated with humoral immune responses and that intra-graft infiltration by ASCs is observed in RAS allografts (*14*).

To investigate the temporal establishment of this intra-graft ASC niche, orthotopic mouse lung transplantations were performed from donor B6D2F1/J mice into B6D2F1/J (isograft), C57BL/6J (RAS) or DBA/2J (BOS) recipient mice (**Fig. 1A**). That humoral immune responses are restricted to RAS phenotype was demonstrated by a progressive increase in circulating donor-specific IgG antibodies over time post-transplantation, a phenomenon absent in both BOS and isograft transplanted mice (**Fig. 1B**). To investigate the temporal evolution of ASC infiltration, histologic sections from RAS lung allografts at days 7, 14, and 28 post- transplantations were examined by immunofluorescence staining for expression of naïve B cell marker B220, plasma cell marker CD138, and IgG to distinguish antibody-secreting capacity (**Fig. 1C**). Temporally, naïve B220^+^ B cells, found in dense clusters within BVBs, peaked at day 7 post-transplant, while CD138^+^IgG^+^B220^−^ ASCs were noted at day 14 and further increased by day 28. The absence of B220 expression in CD138^+^IgG^+^ ASCs suggested that these are terminally differentiated plasma cells. Plasma cell accumulation over time was further quantitated by utilizing *Blimp1^EYFP^* mice utilized as recipients (**Fig. 1D**). Blimp-1, the master transcription regulator of plasma cell differentiation, silences genes expressed in mature B cells and induces genes involved in translational and protein secretion machinery (*30*). FACS analyses of transplanted lung demonstrated CD38^+^B220^low^CD138^+^Blimp-1^+^ population emerge at day 14 and further increase over time **(Fig. 1E),** constituting over 45% of CD45^+^CD3^−^CD4^−^ cells at day 28 post-transplant (**Fig. 1F**). Immunofluorescent analyses of RAS allografts at day 28 post- transplant revealed aggregates of terminally differentiated plasma cells (B220^−^CD138^+^Blimp-1^+^) localized to the BVBs; and negligible numbers of B220^+^Blimp-1^+^ cells were observed in these RAS allografts (**Fig. 1G**).

**Fig. 1.**
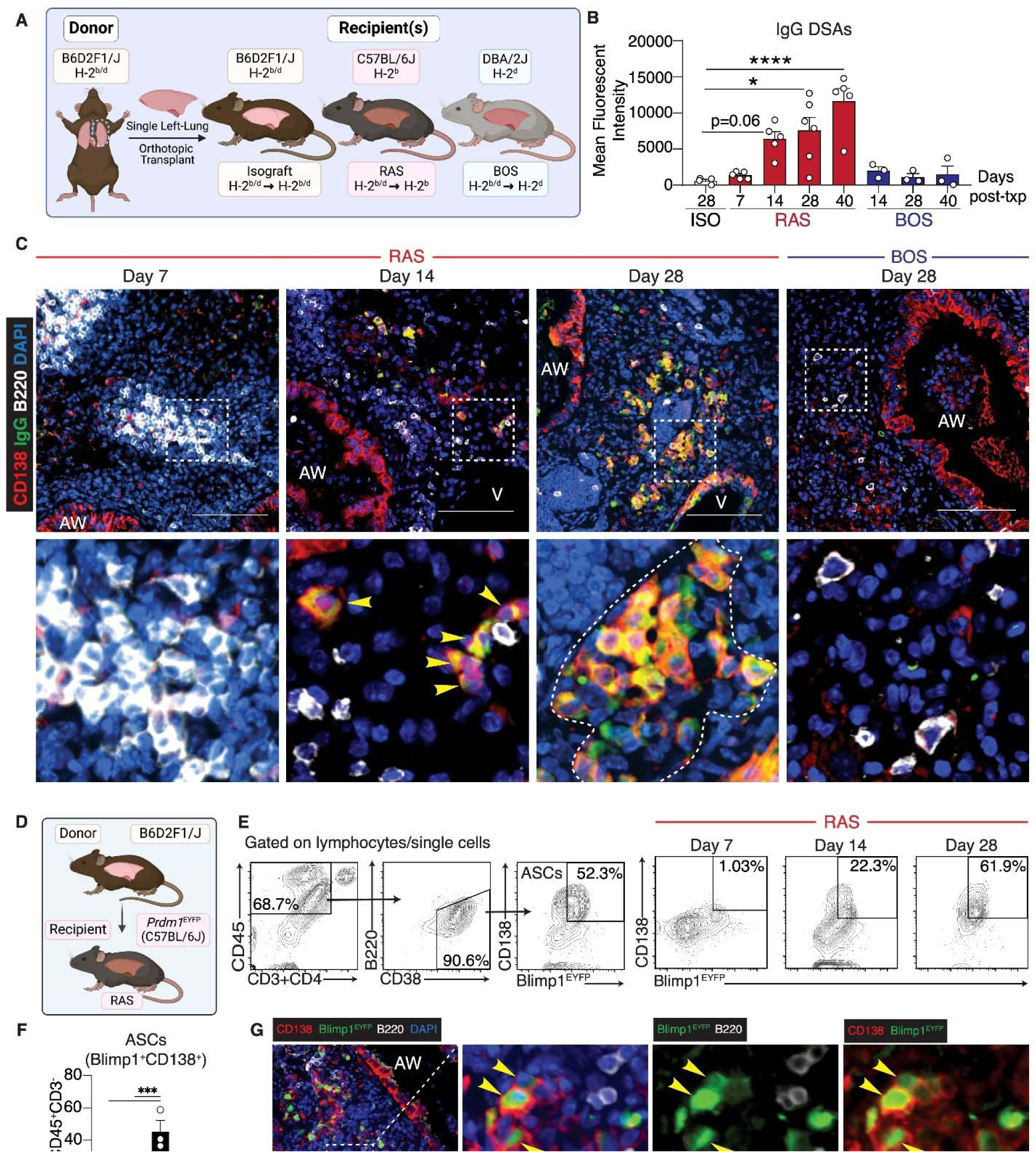
Temporal establishment of pro-survival ASC niches in murine RAS allografts. (**A**) Schemata shows murine models of CLAD for the two phenotypes–RAS (restrictive allograft syndrome), BOS (bronchiolitis obliterans syndrome), and isograft with histocompatibility mismatch. (**B**) Serum alloantibody levels of donor-specific IgG antibodies in isografts, RAS and BOS allografts at the indicated timepoints. n = 3–6 per group. (**C**) Murine RAS and BOS allografts at the indicated timepoints were immunostained for B220, CD138, and IgG. B220 (white) was absent in cells demonstrating co- localization of IgG (green) and CD138 (red) reflecting ASCs. Bottom panels: inset. (**D-G**) *Blimp1^EYFP/+^* C57BL/6J mice were utilized as recipients in the RAS model. (**E**) Characterization of intragraft ASC population (B220^low^CD138^+^Blimp-1^+^) at the indicated timepoints by flow cytometry. CD45^+^CD3^−^CD4^−^ cells were gated for B220^low^CD38^+^ and gated for CD138^+^Blimp-1^+^ subset. (**F**) Percentage of intragraft CD45^+^CD3^−^CD4^−^ ASCs at days 7, 14, and 28. n=3 per group. (**G**) Immunofluorescent staining demonstrating aggregates of terminally differentiated PCs (B220^−^CD138^+^Blimp-1^+^; green) localized to BVBs in day 28 *Blimp1^EYFP^* RAS allografts. Abbreviations: AW, airway; V, vessel. **C, G.** Original magnification: 200x; Scale bar = 50 µm. n=3 per group. **B,F.** Values are represented as means ± SEM. \**p<0.05*, \*\*\**p<0.001*, \*\*\*\**p<0.0001*. One-way ANOVA; post-hoc test: Bonferroni’s.

### Spatial characterization of ASC niche in rejecting lung allografts

Our murine model suggested that ASCs accumulate in rejecting lung allografts of RAS phenotype and that these ASCs localize along the BVBs. To investigate if this is relevant to human CLAD lungs, tissue sections from autopsy or explanted lungs were stained with hematoxylin and eosin (H&E), followed by histologic examination by a blinded pulmonary pathologist for the presence of plasma cells (**Table S1**). Abundant numbers of plasma cells were noted in all the RAS sections (**Table S1**). In contrast, significantly fewer plasma cells were observed in sections from BOS patients even though these end-stage lungs had some mixed features with alveolar damage and pleural fibrosis (**Table S1**).

Anatomically, plasmacytic infiltrates were noted predominantly in two locations: first, along the fibrotic bronchovascular bundles where they were present in close approximation to the scattered mesenchymal cells embedded in the extracellular matrix; second, plasma cells and lymphocytes were noted in the subpleural area in conjunction with fibrotic pleural thickening (**Fig. 2A**). Movat Pentachrome staining for mature collagen and immunofluorescent staining for plasma cell markers CD138^+^CD38^+^IgG^+^ on contiguous slides from human RAS explant lung sections confirmed clusters of plasma cells localized to the leading edge of the advancing fibrosis found interspersed with spindle-shaped MCs (**Fig. 2B**). Semi-quantitative density plot analysis of ASCs and B cells in murine RAS explants stained with Movat Pentachrome and CD138^+^B220^+^IgG^+^ immunofluorescent staining confirmed a clear pattern of plasma cell infiltration along the bronchovascular bundle and in fibrotic subpleural spaces (**Fig. 2C-F**).

**Fig. 2.**
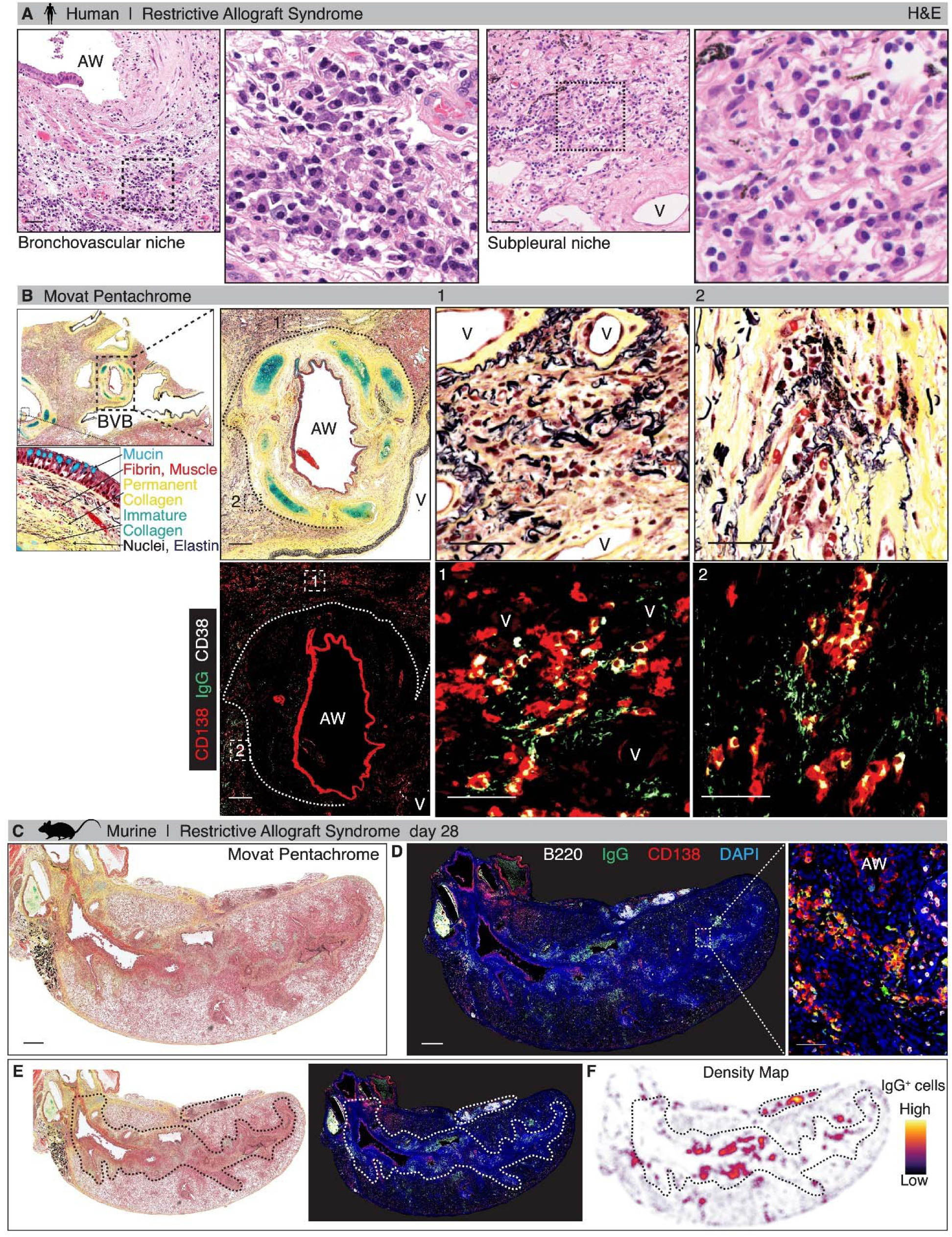
Spatial characterization of plasma cell niche in human and murine RAS lung allografts. (**A**) Representative sections of human RAS lungs. H&E staining highlighting plasma cell clusters in bronchovascular and subpleural niches. (**B**) Consecutive sections from human RAS lungs were stained with Movat Pentachrome to analyze collagen deposition (top panel) and CD138, IgG and CD38 to identify plasma cells (bottom panel). CD138 (red), IgG (green) and CD38 (white) expressing plasma cell infiltrates along the edges of mature collagen in the bronchovascular bundles are shown in insets. n=3, Scale bar = 500 µm; insets = 50 µm. Abbreviations: BVB, bronchovascular bundle; AW, airway; V, vessel. (**C-D**). Representative high-resolution whole slide image of murine RAS allograft characterizing spatial localization of plasma cells. Consecutive sections stained Movat Pentachrome (**C**) and CD138, B220, IgG (**D**) are shown. (**E**) Dotted lines outline clusters of plasma cells, colocalization of CD138 (red) and IgG (green), seen along the bronchovascular bundles and subpleural spaces. (**F**) Density map of IgG^+^ (DAPI^+^) cells is shown marking spatial locations. n=3. Scale bar = 500 µm.

### *Cxcl12^+^ Il6^+^ Gli1^+^ Foxf1^+^* BVB-MCs and the ASC anatomic niche in RAS

CXCL12 is a key cytokine for the establishment of plasma cell niches and regulates key biological processes via ligation of its primary receptor CXCR4 (*31–33*). Our finding of an established ASC niche in the rejecting RAS lung led us to investigate the CXCL12-expressing populations in an adult lung. Among sorted immune (CD45^+^), epithelial (EpCAM^+^), mesenchymal (PDGFRα^+^), and endothelial (CD31^+^) populations, the highest expression of *Cxcl12* was noted in the mesenchymal cells (**Fig. 3A**).

**Fig. 3.**
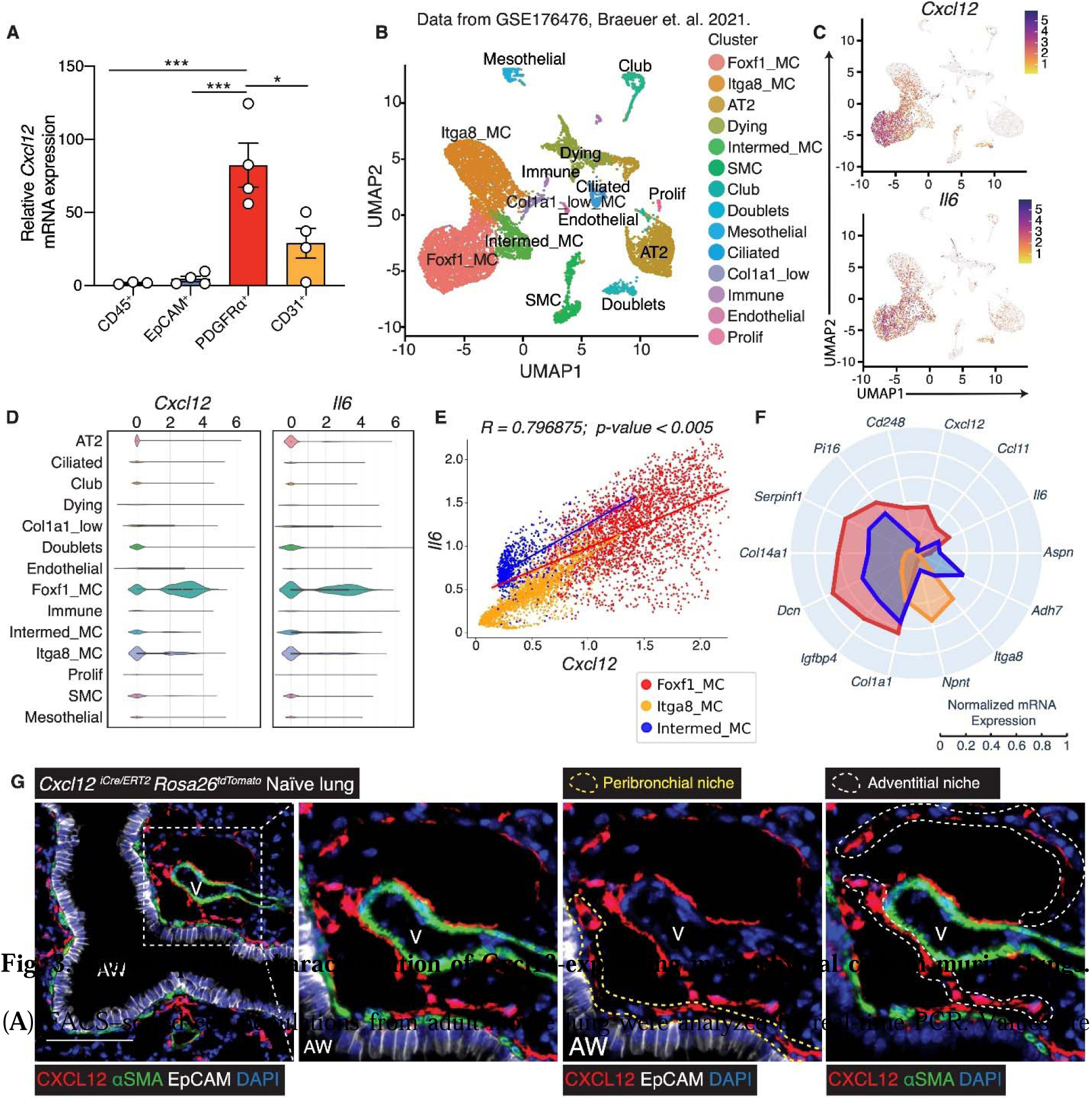
ran cript ic characterization of Cxcl12-expres ng mesenchymal cells in murine lungs. (**A**) FACS–sorted cell populations from adult mouse lung were analyzed by real-time PCR. Values are represented as means ± SEM. \**p<0.05*, \*\*\**p<0.001*. One-way ANOVA; post-hoc test: Bonferroni’s. (**B**) Uniform manifold approximation and projection (UMAP) plot from scRNA-Seq on CD45^−^CD31^−^ cell populations sorted from adult mouse lungs (26). (**C**) UMAP plots of *Cxcl12* and *Il6*, with coordinates same as in **B**, but the color intensity indicates gene expression level within each cell. (**D**) Violin plots show expression levels of *Cxcl12* and *Il6* in each of the 14 clusters. (**E**) Correlation plot showing Foxf1_MC cluster distinctly expressing *Cxcl12* and *Il6* gene expression. (**F**) Radar plot representing differentially expressed key genes marking the signature of each lung mesenchymal clusters. (**G**) *Cxcl12^iCre/ERT2^Rosa26^tdTomato^*adult mouse lung sections were stained for CXCL12 (red), αSMA (green) and EpCAM (white). Dotted lines within insets highlight peri-bronchial (yellow) and peri-vascular adventitial (white) spatial niches for CXCL12^+^ MCs. n=3 per group. Original magnification: 200x; Scale bar = 50 µm. Abbreviations: AW, airway; V, vessel.

Since the site of ASC accumulation in RAS allografts localized with the *in situ* niche of previously described *Gli1^+^Foxf1^+^Itga8^−^Col1a1^+^* BVB-MCs, we plotted *Cxcl12* mRNA expression in the mesenchymal cell clusters previously identified by single-cell sequencing (*26*) (**Fig. 3B-C**). *Cxcl12* expression was found to overlap with the Foxf1_MC cluster (**Fig. 3C-top panel;** **Fig. 3D-left panel**).

IL-6 is a multifunctional cytokine that is considered as a critical component of the PC niche (*34, 35*). The ability of stromal cells to support PC survival is linked to IL-6 (*29, 33, 34, 36, 37*), and IL-6 deficiency can lead to a decreased number of PCs (*34, 36*). *Il6* expression was noted to be unique to the *Foxf1*_MC cluster (**Fig. 3C-bottom panel;** **Fig. 3D-right panel**), with overlap between *Cxcl12* and *Il6* expression (**Fig. 3E**). Other investigators, utilizing single-cell RNA sequencing to characterize resident *Col1a1*^+^ MCs in an adult lung, have also recognized unique subsets associated with BVBs (*38, 39*). Key genes described by other groups as marking specific clusters were plotted across our key clusters using a radar plot as shown in **Fig. 3F**. Foxf1_MC cluster expressed the highest levels of key genes shown to be expressed in BVB-associated MCs by Tsukui et al (*38*). These included genes marking *Col1a1*^+^ MC in the peri-vascular adventitial cuffs (*Pi16* and *Ccl11*) and peri-bronchial region (*Dcn*). *Adh7* expression, which has been previously demonstrated in non-cytokine producing adventitial cells in the BVBs, was absent in the Foxf1_MC cluster but was expressed in the intermediate cluster (**Fig. 3F**) (*38*). The Itga8_MC cluster, which represents the alveolar cluster in our data set, expressed *Npnt* which was shown to be a marker for *Col1a1*^+^ cells in the alveoli (*38*).

To further investigate the CXCL12-expressing MCs, tamoxifen-dosed adult *Cxcl12^iCre/ERT2^Rosa26^tdTomato^* mice were utilized. Real-time PCR confirmed high *Cxcl12, Il6, Dcn, Igfbp4, Col14a1, Gli1* in flow sorted PDGFRα^+^ tdTomato-CXCL12^+^ population (**Fig. S1A-B**). On immunofluorescent staining of lungs harvested from *Cxcl12^iCre/ERT2^Rosa26^tdTomato^*, CXCL12- related-tdTomato expression was restricted to the MCs in BVBs (**Fig. 3G**) and pleural space (**Fig. S1C**). Within BVBs, intrinsic expression of CXCL12 was noted in sub-bronchial MCs adjacent to epithelial cells, bronchial cuff (outside smooth muscle cells), and the peri-vascular adventitial cuff cells (**Fig. 3G**). This expression pattern overlaps with that of *Foxf1* noted previously in our *Foxf1^tdTomato^*adult mice (*26*).

Next, to investigate the expression of *Cxcl12* and *Il6* in the allograft milieu, sorted cell populations from day 28 allografts were compared to isografts for the expression of these cytokines (**Fig. 4A**). Significantly higher *Cxcl12* and *Il6 mRNA* expression was noted in the PDGFRα^+^ MC populations in allografts compared to isograft controls, while no induction was noted in the other cell types (**Fig. 4A**). To further assess the role of CXCL12^+^ MCs in the establishment of ASC niches in RAS pathogenesis, tamoxifen-dosed *Cxcl12^iCre/ERT2^Rosa26*^tdTomato^ B6D2F1/J donor mice were transplanted into wildtype C57BL/6J mice (**Fig. 4B**). Immunofluorescent staining of RAS lung allografts at day 28 post-transplantation demonstrated expansion of the CXCL12^+^ cells in the BVBs (**Fig. 4C**). Aggregates of CD138^+^IgG^+^ ASCs were noted lying in close approximation to these tdTomato- CXCL12^+^ MCs, lying in the same anatomic location and at the fibrotic pleura (**Fig. 4D, Fig. S1C-D**). These data demonstrate that in the rejecting lung allograft, there is an expansion of CXCL12-expressing MC populations along with upregulation of *Cxcl12* and *Il6* expression.

**Fig. 4.**
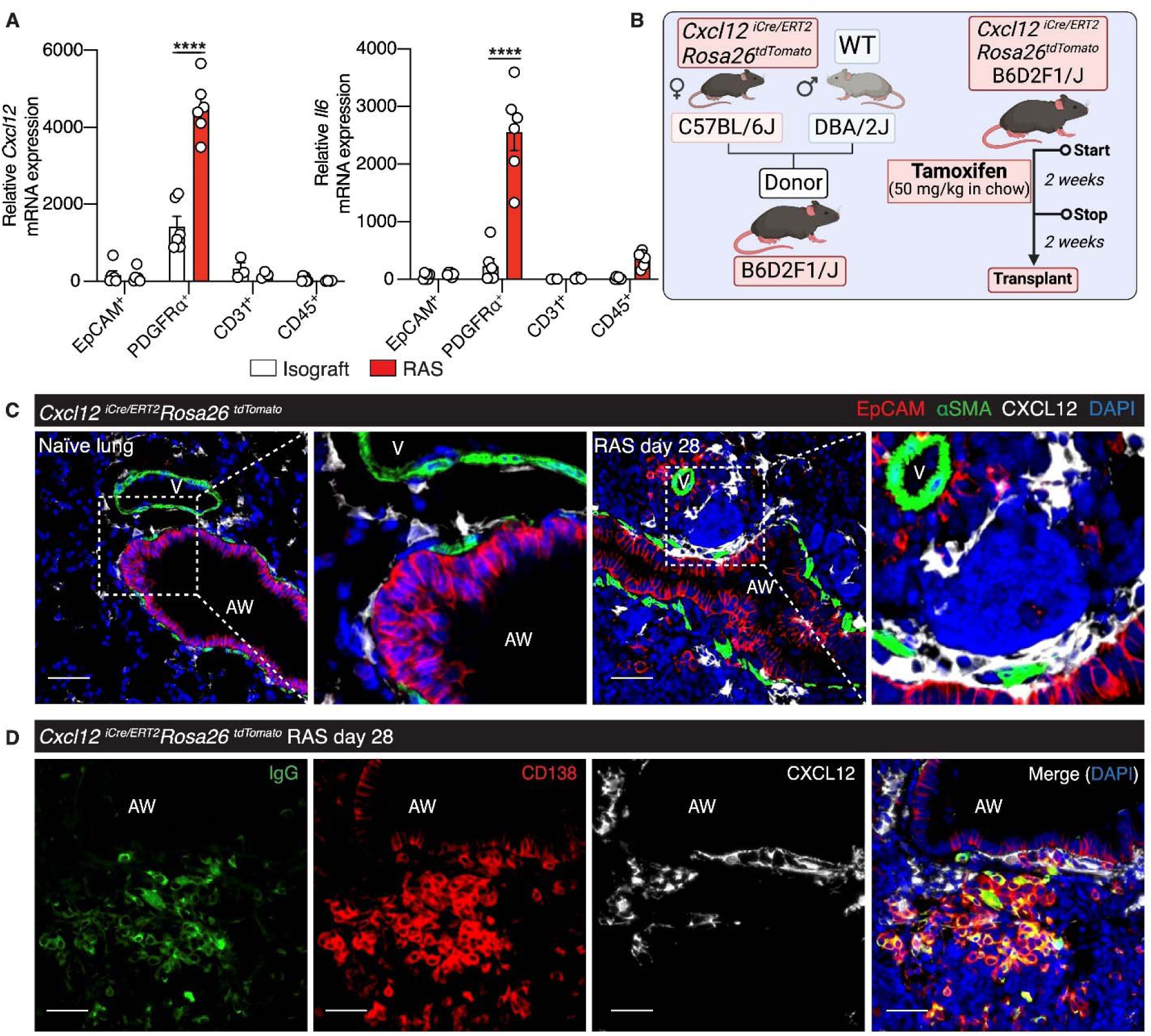
Increased CXCL12 expression and expansion of CXCL12+ MCs provide for an ASC niche in RAS lung allografts. (**A**) *Cxcl12* and *Il6* mRNA expression analyzed by real-time PCR in FACS-sorted cell populations from an isograft and day 28 RAS allografts. Values are represented as means ± SEM. ****, *p<0.0001*. One-way ANOVA; post-hoc test: Bonferroni’s. (**B**) Schema for the generation of *Cxcl12^iCre/ERT2^Rosa26^tdTomato^*B6D2F1/J donor mice and lung transplant experiments. (**C**) Representative immunofluorescent staining of *Cxcl12^iCre/ERT2^Rosa26^tdTomato^*naïve and transplanted lungs stained for CXCL12 (white) αSMA (green) and EpCAM (red). (**D**) Representative immunofluorescent staining for CD138 (red) and IgG (green) in *Cxcl12^iCre/ERT2^Rosa26^tdTomato^*donor mice demonstrating plasma cells colocalizing with CXCL12^+^ MCs. n=3 RAS allografts. Scale bars: 50 μm.

### IL-6 trans-signaling as a novel regulator of CXCL12 in human lung-resident MCs

Our finding that *Cxcl12* is upregulated in PDGFRα^+^ cells isolated from murine RAS allografts led us to investigate regulators of CXCL12 expression in MCs. MCs derived from human lung transplant recipients with RAS demonstrated significant *CXCL12* mRNA expression as compared to normal non-CLAD controls (**Fig. 5A**). Treatment with key pro-fibrotic agents, transforming growth factor-β (TGF-β) and lipid mediator lysophosphatidic acid (LPA) did not affect CXCL12 expression in non-CLAD MCs (**Fig. 5A**). We have previously demonstrated that human lung MCs express IL-6 but lack IL-6 receptors, further finding that IL-6 trans-signaling is active in these cells and leads to their fibrotic differentiation (*40*). Significant upregulation in

**Fig. 5.**
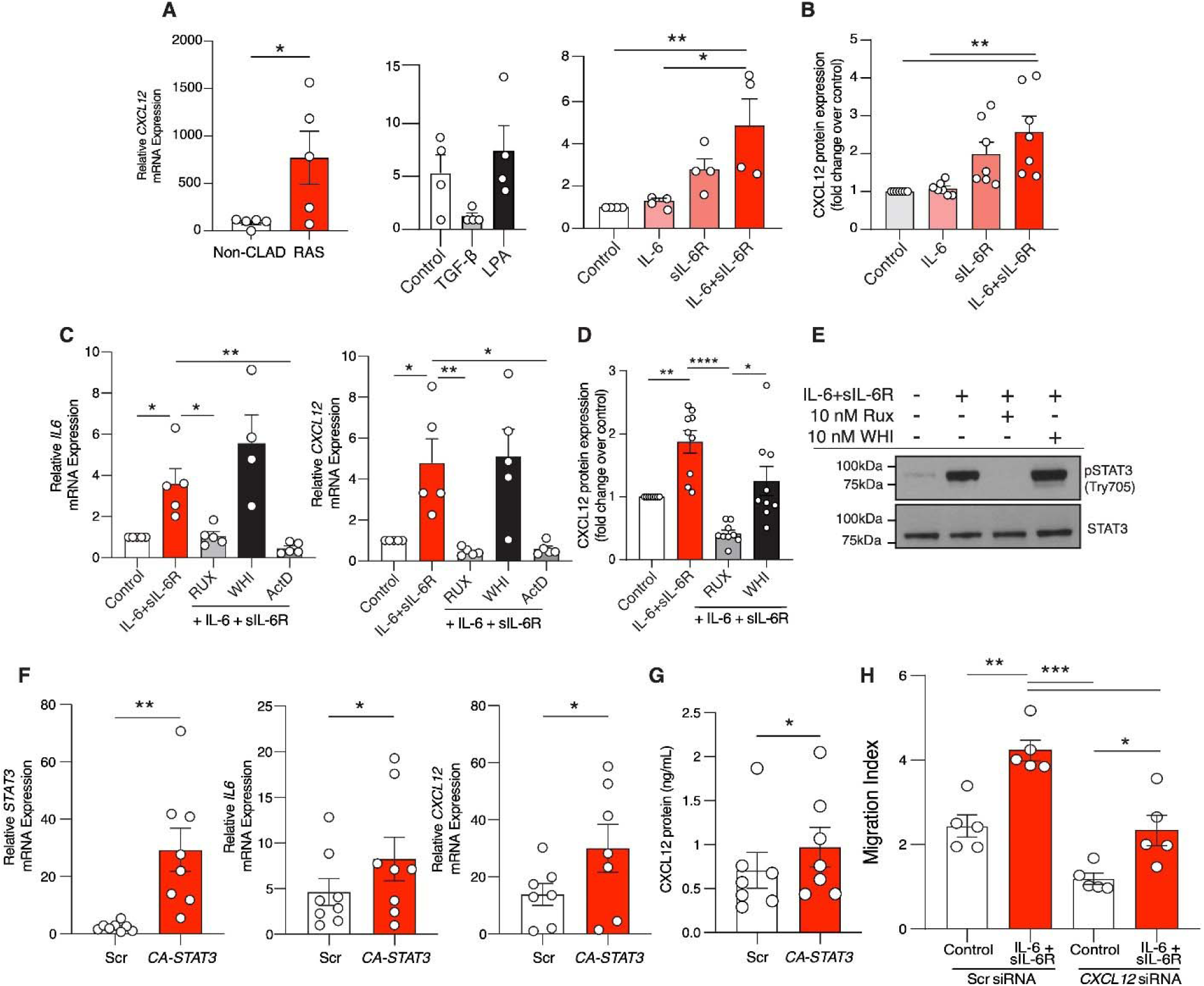
IL-6 trans-signaling induces CXCL12 expression in human MCs. (**A**) *CXCL12* mRNA expression analyzed by real-time PCR in MCs from non-CLAD lung transplant patients or diagnosed with RAS; in non-CLAD MCs treated with TGF-β or LPA; and IL-6, sIL-6R or both. (**B**) CXCL12 protein expression in conditioned media (ELISA) from non-CLAD MCs treated with IL-6, sIL-6R or both. (**C-E**) Non-CLAD MCs pre-treated with JAK1/2 pan-inhibitor, Ruxolitinib (Rux), or JAK3 inhibitor, WHI-P154 (WHI), and pan-transcriptional inhibitor, Actinomycin D (ActD), were stimulated with IL-6 and sIL-6R. *IL6, CXCL12* mRNA expression (real-time PCR) and CXCL12 protein expression (ELISA) are shown in C and D, respectively. (**E**) Representative western blot of STAT3 phosphorylation (Tyr705). (**F,G**) Non-CLAD MCs were transfected with lentiviral vectors that allowed constitutional activation of STAT3 (CA-STAT3) or scrambled controls. (**F**) *STAT3, IL6, CXCL12* mRNA expression (real-time PCR). (**G**) CXCL12 protein expression in conditioned media (ELISA). (H) Non-CLAD MCs were transfected with scrambled control or *CXCL12* sequence specific siRNAs, and then stimulated with IL-6 and sIL-6R (48h). Conditioned media was used to stimulate 0.5 million splenocytes in the upper chamber to migrate through the membrane. Total CD45^+^CD3^−^CD19^+^ splenocytes in the lower chamber was measured by FACS analyses. Values = means ± SEM. \**p<0.05*, \*\**p<0.01*, \*\*\**p<0.001*, \*\*\*\**p<0.0001*. Student’s t-test (**A,F-G**). One-way ANOVA; post-hoc test: Bonferroni’s (**B-D,H**).

CXCL12 at both mRNA and protein levels was noted in non-CLAD MCs treated with a combination of IL-6 and sIL-6R (**Fig. 5A,B**). Interestingly, we also noted an increase in *IL6* mRNA expression suggesting that IL-6 trans-signaling further induces *IL6* expression in these MCs (**Fig. 5C**). This increased *CXCL12* and *IL6* mRNA expression with IL-6 trans-signaling noted to be abrogated by JAK1/2 inhibitor Ruxolitinib and pan-transcriptional inhibitor actinomycin D, but not JAK3-specific inhibitor WHI-P154 suggesting JAK1/2-STAT3 transcriptional regulation (**Fig. 5C**). CXCL12 secretion was assessed by ELISA in conditioned media from these conditions and demonstrated a similar trend (**Fig. 5D**). Confirmatory immunoblotting analyses demonstrated that inhibition of JAK1/2, but not JAK3, effectively blocked IL-6 trans-signaling-mediated STAT3 phosphorylation (**Fig. 5E**). These findings suggest an obligatory role for IL-6 trans-signaling-induced STAT3 phosphorylation (Tyr705) in the upregulation of CXCL12 expression.

To confirm STAT3-mediated transcriptional regulation of CXCL12, CLAD-free MCs were transduced with lentiviral particles containing constitutively active STAT3 vectors (*CA- STAT3*) or empty backbone control (*Scr*). Transduction efficiency was confirmed by analyzing the mRNA expressions of *STAT3* and *IL6* (**Fig. 5F**). Real-time PCR analyses of *CXCL12* mRNA expression demonstrated 10-fold higher levels in the MCs expressing constitutively active STAT3 compared to controls, which was confirmed by ELISA on conditioned media (**Fig. 5F- G**). To determine the chemoattractant potential of IL-6 trans-signaling-induced mesenchymal CXCL12 expression, we assessed the migratory capacity of primary lymphocytes in response to *CXCL12*-silenced MCs by FACS analyses. CD45^+^ splenocytes from naïve adult C57BL/6J mice cultured in the upper chamber of transwell inserts were exposed to conditioned media from *CXCL12-*silenced CLAD-free MCs or scrambled controls in the presence or absence of IL-6 trans-signaling. Compared to unstimulated scrambled controls, stimulation with IL-6 trans-signaling induced ∼2-fold higher migration of splenocytes, abrogated by *CXCL12-*silencing (**Fig. 5H**).

### IL-6 trans-signaling blocker Olamkicept inhibits ASC accumulation and ameliorates fibrosis in murine RAS allografts

To further investigate the *in vivo* relevance of this proposed IL-6 trans-signaling/CXCL12 pathway in the establishment of local ASC niche and allograft fibrogenesis, we targeted IL-6 trans-signaling in our orthotopic murine lung transplant model of RAS utilizing a pharmacological approach. Lung allograft recipients were treated with the trans-signaling blocker Olamkicept or saline control administered once weekly by intraperitoneal injection starting on day 3 (**Fig. 6A**). Olamkicept-treated RAS allografts demonstrated ∼50% lower levels of CXCL12 protein expression in lung homogenates compared to saline-treated allograft controls (**Fig. 6B**). Compared to RAS allograft controls, Olamkicept treatment reduced circulating donor- specific IgG by ∼40%, a decrease comparable to that of proteasome inhibitor Bortezomib (**Fig. 6C**), although circulating donor-specific IgMs were unaffected (**Fig. 6D**). To further determine if Olamkicept treatment can inhibit ASC accumulation in the grafts, we utilized *Blimp1^EYFP/+^*reporter mice as recipients and assessed infiltrating immune populations by FACS analysis in the presence of Olamkicept or saline control treatments. Olamkicept not only decreased total Blimp- 1^+^CD138^+^ ASCs (**Fig. 6E**), but also decreased total CD45^+^ cells, CD45^+^CD3^−^CD4^−^ Blimp-1^+^ cells, and CD19^+^ cells (**Fig. 6E**). Immunostaining analyses of FFPE sections confirmed a decrease in BVB infiltration by ASCs (CD138^+^IgG^+^), further confirming a role for IL-6 trans-signaling in BVB-MC-mediated ASC recruitment to the allograft (**Fig. 6F**). Immunohistochemical staining to detect CCSP (club-cell secretory protein) expression was performed to investigate if Olamkicept protects airway epithelial integrity targeted by humoral responses in RAS allografts (*14*). While day 28 control allografts demonstrated almost complete loss of airway CCSP staining as previously described, CCSP expression was preserved in Olamkicept-treated allografts (**Fig. 6F**). Movat Pentachrome staining on day 28 RAS allografts treated with Olamkicept presented decreased fibrosis around bronchovascular bundles, indicated by a reduction in permanent collagen (yellow) stained regions, compared to allograft controls (**Fig. 6F**). A quantitative assessment of collagen deposition utilizing hydroxyproline assay demonstrated a significant reduction in Olamkicept-treated allografts compared to controls (**Fig. 6G**).

**Fig. 6.**
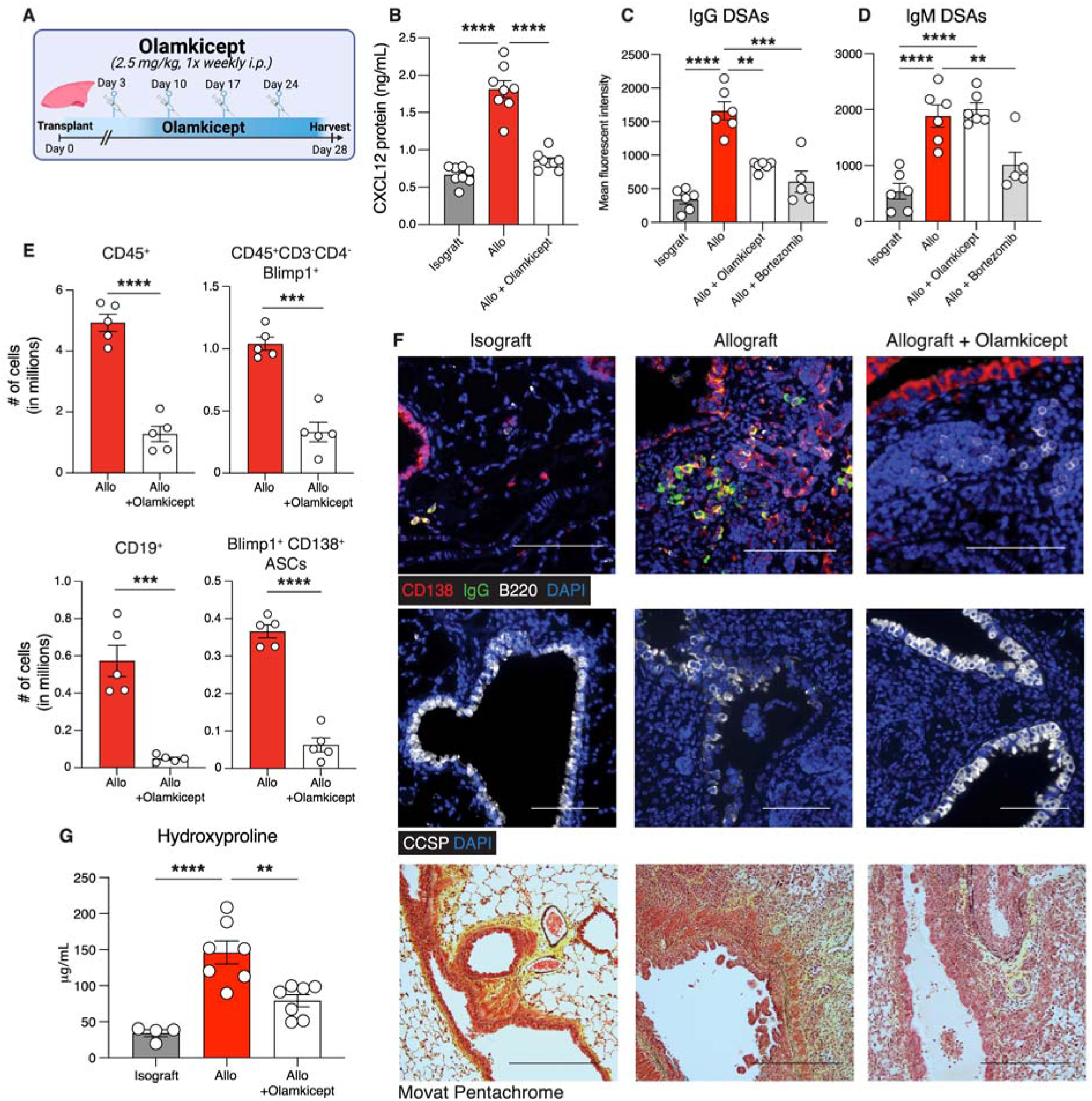
IL-6 trans-signaling neutralization by Olamkicept suppresses humoral immune responses and fibrosis in RAS lung allografts. (**A**) Experimental schematic. B6D2F1/J donor lungs were transplanted into C57BL/6J or *Blimp1^EYFP^* C57BL/6J recipient mice, followed by treatment with Olamkicept (2.5 mg/kg/week; intraperitoneal injections) between days 3 and 28. (**B**) CXCL12 protein concentration by ELISA in lung homogenates from **A**. (**C,D**) Isografts, or RAS allografts (Allo) treated with or without Olamkicept or Bortezomib (2.5 mg/kg) were sacrificed at day 28. Serum was analyzed for donor-specific IgG (**C**) and IgM (**D**) levels by flow cytometry. (**E**) FACS analyses of single cell suspensions of allografts from *Blimp1^EYFP^* C57BL/6J recipients treated with Olamkicept. (**F**) Immunofluorescent staining of representative tissue sections showing naïve B cells (B220, white) and antibody-secreting cells (CD138, red; IgG, green), top panels; airway epithelial integrity, marked by CCSP (white), middle panels; and stained for Movat Pentachrome, bottom panels. n=3. Original magnification = 200x. Scale bar = 50 µm. (**G**) Quantitative assessment of fibrosis by measuring hydroxyproline content in graft lung homogenates. Values are represented as means ± SEM. \*\**p<0.01*, \*\*\**p<0.001*, \*\*\*\**p<0.0001*. Student’s unpaired t-test (**E**). One-way ANOVA; post-hoc test: Bonferroni’s (**B-D,G**).

## DISCUSSION

In this study, we investigate mesenchymal-immune cell interactions in rejecting lung allografts and elucidate the role of specialized stromal cells in establishing a stable intra-graft survival niche for antibody-secreting cells (ASCs). We identify a CXCL12 and IL-6 expressing stromal cell population restricted along the bronchovascular bundles, comprising the Foxf1^+^ population previously implicated in allograft fibrogenesis (*26*). A novel IL-6 trans-signaling/JAK STAT-dependent mechanism of CXCL12 regulation in mesenchymal stromal cells was identified. Notably, we demonstrate that pharmacologic inhibition of IL-6 trans-signaling prevents the formation of intra-graft ASC niche and ameliorates fibrosis in a murine lung transplant model. Thus, we identify a novel paradigm of targetable paracrine IL-6 trans- signaling/CXCL12 signaling axis between graft-resident mesenchymal stromal cells and humoral immune cells in regulating the inflammatory niche and graft fibrosis in rejecting lungs.

Our studies, utilizing human lung allografts and novel murine orthotopic lung transplant models of chronic allograft rejection, provide translational insights into the spatiotemporal establishment of pro-survival ASC niches in CLAD pathogenesis. ASC accumulation and its contribution to disease via local antibody secretion have been described in various inflammatory/immune conditions such as lupus nephritis (*41, 42*) and chronic rhinosinusitis (*43, 44*). Examination of human explanted RAS lungs demonstrated clusters of plasma cells, identifiable on plain histology and confirmed by CD138 staining. Our comparison of RAS and BOS human lungs confirmed significantly higher ASC accumulation in RAS. However, patients can show mixed features of both phenotypes, and some plasma cells have been observed in BOS patients (*4, 45, 46*). ASCs comprise plasmablasts and fully differentiated PCs, with long-lived PCs demonstrating resistance to specific immunosuppressive therapies (*33*). Immunoglobulin G and CD138 expression did not overlap with B220 expression in murine RAS lung allografts, suggesting these are fully differentiated PCs, a finding corroborated by using *Blimp1^EYFP^*mice as recipients in the murine RAS lung transplant model. A striking finding was the specific anatomic localization of these ASCs along the BVBs and in the subpleural space in both human and murine grafts, providing evidence for a specific pro-survival niche and interaction of infiltrating immune cells with graft-resident MCs.

We identify the anatomically and functionally distinct subset of MCs that express CXCL12 in an adult lung and support the ASC niche. The key role of specialized stromal cells in the establishment of a stable survival niche for ASCs is well-recognized across a variety of lymphoid and hematopoietic tissues (*27–29*). CXCL12 gradient produced by specific medullary fibroblastic reticular cells, direct the migration of plasmablasts to the medullary cords of the lymph nodes to become mature plasma cells. These anatomically, phenotypically, and functionally distinct subset of mesenchymal stromal cells in the lymph node are also a local source of plasma cell survival factors that include IL-6, BAFF, and APRIL (*27*). Similarly, it is well established that the migration of circulating plasmablasts across the endothelium into the specialized microenvironments in the bone marrow parenchyma is directed by CXCL12. After arrival of motile plasmablasts, they become sessile, docking onto, and remaining in close contact with the stromal cells. This CXCL12-expressing sub-population of stromal cells within the bone marrow is also required for the survival of mature long lived plasma cells in the bone marrow (*47, 48*). We found ASCs localized along the BVB and that previously described Gli1^+^Foxf1^+^ MCs, which reside in the same anatomic niche, uniquely express CXCL12 and IL-6. Characterization of CXCL12-expressing population in adult lungs by utilizing CXCL12 *Cre* mice confirmed that CXCL12 expression is limited to cells along the BVBs. Utilizing the lineage-traced *Cxcl12 Cre* mice as donors in our murine lung transplant model, we confirmed their expansion and close approximation to ASCs in a rejecting allograft. Our findings that this cell population is collaborated by a recent report identified IL-33 expressing adventitial stromal cells in BVBs as key for type 2 lymphocytic expansion and function (*39*). Further understanding of the subpopulations within these BVBs and the factors influencing the gradient of chemokines and cellular migration, and homing are required.

Another striking finding was the demonstration that IL-6 trans-signaling augments CXCL12 expression in MCs. Human lung-resident MCs upregulated CXCL12 expression in response to IL-6 and sIL-6R via the JAK1/2-STAT3 signaling pathway. The *in vivo* relevance of this pathway was established in the murine RAS model, where targeting IL-6 trans-signaling led to decreased CXCL12 expression. Our previous work evaluated IL-6 signaling in the context of lung transplantation and CLAD, noting increased IL-6 and IL-6 receptor expression in BAL samples from CLAD patients (*40*). IL-6 signaling occurs via two pathways: classical (cis- signaling) and trans-signaling. Classical signaling involves IL-6 binding to the IL-6 receptor (IL- 6R) and glycoprotein 130 (gp130) heterodimer, activating the JAK/STAT pathway. IL-6R is expressed on a limited subset of cells (*49*). In contrast, trans-signaling occurs when soluble IL- 6R (sIL-6R), produced by alternative mRNA splicing or proteolytic cleavage, binds IL-6 extracellularly (*50–53*). Our published work highlighted crosstalk between lung MCs and mononuclear cells, where MCs induce mononuclear cells to secrete more IL-6 and shed sIL-6R, which in turn induces vigorous IL-6 trans-signaling in lung MCs, stimulating JAK2-dependent STAT3 phosphorylation and nuclear translocation (*40*). Macrophage infiltration is an early and prominent feature of RAS allografts (*14*). The interaction of CXCL12-expressing stromal cells and myeloid cells is crucial for supporting ASC niches, but the underlying mechanisms are not fully understood. Our data suggest a novel signaling paradigm where sIL-6R secreted by mononuclear cells triggers IL-6 trans-signaling in MCs, leading to CXCL12 induction and ASC niche establishment.

The role of the IL-6 trans-signaling/CXCL12 induction in regulating local humoral immune responses in the lung has not been previously recognized, and we believe these are the first studies to investigate this signaling pathway in CLAD. RAS is a particularly aggressive CLAD phenotype with a median survival of less than one year and few effective therapeutic options (*7*). The recognition of its link to AMR and increased DSAs has led to the use of modalities aimed at targeting HLA antibodies, such as plasmapheresis, intravenous immunoglobulins, and rituximab (*7–9, 11–13, 54, 55*). While partially successful at reducing the serum antibody titers, these approaches have not proven efficacious in treating or stabilizing RAS (*55*). Resistance to immunosuppressants and B-cell depletion therapies like rituximab in other inflammatory and autoimmune diseases has been linked to the survival of long-lived PCs, which downregulate B cell markers like CD20 and can evade clearance (*56–59*). Our findings support a similar mechanism in murine RAS lungs, where we demonstrate CD138^+^IgG^+^B220^−^ ASCs in BVB niches. Proteasome inhibitor Bortezomib has been shown to effectively deplete both short- and long-lived plasma cells in murine models (*60*), and investigations have also utilized concurrent ASC and B cell-depleting therapies (*61, 62*). Our findings of the critical role of IL-6/sIL-6R trans-signaling in CXCL12 induction in lung MSCs and maintenance of the pro-survival niche of ASCs in the graft provide a novel rationale for targeting this pathway in RAS. Currently, there are several FDA-approved medications designed to target the IL-6 signaling pathway, including trans-signaling blocker (Olamkicept), neutralizing antibodies against IL-6 (Sirukumab, Olokizumab, Siltuximab), and IL-6R (Tocilizumab, Sarilumab), and JAK inhibitors (Tofacitinib, Ruxolitinib) (*50, 63*). We provide the first evidence of trans the trans-signaling inhibitor Olamkicept in both ameliorating local humoral immune cell infiltration and fibrosis in a well- established murine model of RAS. Interestingly, along with decreased PCs in the graft, we found a decrease in circulating IgGs, suggesting that the trans-signaling inhibition could also be affecting systemic humoral immune response. This decrease in IgGs was comparable to Bortezomib, suggesting a rather significant contribution of IL-6 trans-signaling to PC survival and function. The reduction in collagen and graft fibrosis with Olamkicept was significant but can likely be further improved by combination therapies targeting both humoral and cellular immune responses, which will be the focus of future investigation. Importantly, as therapies targeting IL-6/IL-6R trans-signaling are already in clinical use, these studies can be translated rapidly to meaningfully impact patient care and outcomes after lung transplantation.

Despite the promising results in a disease relevant murine lung transplant model and use of human cells to elucidate mechanistic pathways, there are limitations to clinical translation of our study. First and foremost, we focus on RAS, a specific CLAD phenotype, utilizing a murine model with demonstrated humoral immune response dependency (*64*). However, CLAD phenotypes are determined clinically and there can be significant overlap among various phenotypes (*4*). Furthermore, diagnosis of antibody mediated rejection is difficult and better markers of CLAD endotypes are needed for designing clinical trials which can test endotype guided therapies (*46, 65*). Second, our murine model utilizes F1 to parent mouse transplantation without immunosuppressive medications which are utilized in clinical post-transplant management. Investigating combination therapies in preclinical models or considering the addictive effect of IL-6 trans-signaling inhibition to standard treatment regimens is required.

In summary, we characterize the CXCL12-expressing MCs that provide for a pro survival niche for ASCs in a rejecting lung allograft and specifically demonstrate the contribution of IL-6 trans-signaling to CXCL12 expression, humoral immune niche establishment, and allograft fibrosis. Our investigations open a new chapter into understanding of factors regulating the establishment of intra-graft immune cell niches after solid organ transplantation. Importantly, we demonstrate that characterizing signals, such as IL-6 trans-signaling, which require complex interactions between infiltrating immune cells and graft-resident stromal cells (*40*), and then downstream key paracrine actions of these stromal cells can be harnessed to ameliorate chronic allograft rejection.

## MATERIALS AND METHODS

### Study Design

In this study, we characterize a distinct CXCL12^+^ subpopulation of MCs that support a pro-survival niche for ASCs in rejecting lung allografts, determine regulation of CXCL12 in patient-derived MCs, and assess the efficacy of pharmacologic neutralization of IL-6 trans-signaling to attenuate CXCL12-mediated ASC niche *in vivo*. To accomplish this, we analyzed multiple murine models of CLAD, and retrospective normal, RAS, and BOS human lung tissue sections, for the presence of ASCs. Mice were between 8-16 weeks of age at time of transplant, and both male and female mice were used. For treatments with Olamkicept, Bortezomib, or respective controls, mice were randomly assigned to treatment groups. Power analysis was performed with G Power 3.1, using historical means, standard deviations from our reported experiments to calculate effect parameters using a power of 80%, an α of 0.05 to detect a difference, accumulating a minimum of these samples for any end point. All murine experiments were performed under an approved IACUC protocols at the University of Michigan and Emory University. Histological review of human lung tissue was conducted in a blinded fashion, and tissue sections and lung transplant patient-derived mesenchymal cells were obtained under IRB approval #HUM00042443. RAS patients were diagnosed using International Society for Heart and Lung Transplantation (ISHLT) guidelines (*4*), MCs derived from these samples were labeled RAS. MCs from CLAD-free patients without evidence of rejection, infection, and remained CLAD-free for ≥ one-year post-BAL were labeled non-CLAD-MCs. All data were included and reported without excluding outliers. Clinical data regarding patients in this study are presented in **Table 1**.

**Table 1.** Patient demographics. This table reports the gender, age, specific pre-transplant pathologies, CLAD status, and time post-transplant statistics of each patient derived mesenchymal cell line used in the present study.

| Variable | Entire Cohort (n=23) |
| --- | --- |
| Age (years ± SD): | 47 ± 13.93 |
| Age in years (n, %): |  |
| <30 | 3, 13.04% |
| 30-40 | 4, 17.39% |
| 40-50 | 2, 8.70% |
| 50-60 | 12, 47.83% |
| >60 | 2, 8.70% |
| Female sex (n, %): | 6, 26.09% |
| Pre-transplant diagnosis (n, %): |  |
| COPD/emphysema | 5, 21.74% |
| Cystic Fibrosis | 5, 21.74% |
| ILD | 11, 47.83% |
| Other | 2, 8.70% |
| CLAD status at BAL (n, %): | 6, 26.09% |
| Days post-transplant to CLAD diagnosis (mean ± SD) | 1116 ± 716 |
| Days from CLAD diagnosis to BAL (mean ± SD) | 183 ± 168 |

**Footnote:** CLAD, chronic lung allograft dysfunction; n, number; SD, standard deviation; COPD, chronic obstructive pulmonary disease; ILD, interstitial lung disease; BAL, bronchoalveolar lavage.

#### Sex as a biological variable

Mesenchymal cells were cultured from lavage fluid derived from lung transplant patients of both female and male sex (**Table 1**). Male and female mice were utilized in mouse lung transplant experiments. None of the experiments were limited to samples from either sex. Therefore, sex was not evaluated as a biological variable in the experiments.

#### Study approval

All participants provided written, informed consent prior to participation in the study in compliance with the Helsinki declaration. The study was conducted in accordance with relevant guidelines and regulations using a protocol for human studies approved by the Institutional Review Board at the University of Michigan and the Emory University (approval #HUM00042443). All murine experiments were performed with the approval of the Institutional Animal Care and Use Committee and the Institutional Review Board at the University of Michigan and the Emory University.

#### Mice and orthotopic lung transplant model

Specific pathogen-free male mice B6D2F1/J, C57BL/6J, and DBA/2J were purchased from the Jackson Laboratory, Bar Harbor, ME, and the transplant donors and recipients and breeding colonies were maintained at Emory University. Mice used for transplants were at least 8-16 weeks and weighed between 24-32g. Isograft transplants were performed using a B6D2F1/J to B6D2F1/J strain combination (*14, 64*), and allogenic transplants were performed using an F1 to parent: B6D2F1/J lungs to C57BL/6J combination for the RAS model (*14*), and B6D2F1/J lungs to DBA/2J for the BOS model (*64*). Orthotopic left lung transplantations were performed as previously described (*14, 64*) using a surgical microscope (SZX16-SZX2; Olympus) with 2.1× to 34.5× magnifications for all procedures. Buprenorphine was given to recipient mice at the conclusion of the procedure and every 12 hours until 3 days after transplant. No immunosuppressive drugs were used.

#### Transgenic mice

To label plasma cells, B6.Cg-Tg(*Prdm1^EYFP^*)1Mnz/J (JAX, Stock#:008828) or *Blimp1^EYFP^* mice on a C57BL/6J background were used as transplant recipients with B6D2F1/J donor lungs to label plasmablasts and plasma cells in the RAS model. *Gli1^CreERT2/WT;^Rosa26^mTmG/WT^*mice were generated as previously described (*26*). Briefly, *Gli1^CreERT2/WT;^Rosa26^EYFP^* mice were generated by crossing *Gli1^CreERT2^* with B6.129X1- *Gt(ROSA)26Sortm1(EYFP)Cos/J* (JAX, Stock#006148). The *Gli1^CreERT2/WT^;Rosa26^mTmG/WT^*and *CXCL12^iCre/ERT2/WT^;Gt(ROSA)26^Sortm4(ACTB-tdTomato,-ERFP)^*. B6D2F1/J donor mice were dosed with tamoxifen chow for 14 days at 4-6 weeks of age. Mice were given normal chow for 3 days and then transplanted into C57BL/6J recipients.

#### Statistical Analysis

When comparing the means of two groups, the Student’s two-tailed t test was used to determine “p” values. When comparing the means of three or more groups, one-way analysis of variance was performed with a post hoc Bonferroni test to determine which groups showed significant differences unless otherwise specified. A “p” value of less than 0.05 was considered significant and was analyzed using GraphPad Prism (ver.8.0.0) for Windows 64-bit.

## List of Supplementary Materials

Fig. S1 Table S1

Materials and Methods – Additional Information

## Acknowledgments

We thank the experienced personnel at the Flow Cytometry Core, Research Histology Core (NIH P30 CA04659229), Microscopy Core, Advanced Genomics Core, Bioinformatics Core, and Transgenic Animal Model Core at the University of Michigan for their contributions to this study. We thank the experienced personnel at the Flow Cytometry Core, Research Histology Core, Microscopy Core, and the Emory Integrated Genomics Core at the Emory University for their contributions to this study. The schematic figures were generated on BioRender.

## Funding

National Institutes of Health – NHLBI – R01HL162171, HL094622 (VNL)

The McKelvey Transplant Center (VNL)

The Cystic Fibrosis Foundation LAMA16XX0 (VNL)

The Campbell Gift Fund

## Author contributions

Conceptualization: VNL, APM, NMW, KT Methodology: APM, KT, NMW, RV, YI Investigation: APM, KT, NMW, RV, YI, AR, MM Visualization: VNL, APM, RV, YI, KT

Pathology review: CFF, VNL Funding acquisition: VNL Project administration: VNL Supervision: VNL

Writing – original draft: VNL, APM, RV, KT Writing – review & editing: VNL, APM, RV, KT

## Data and materials availability

Accession numbers for data presented in **Fig. 3B** was previously deposited in NIH GEO MIAME: GSE176476 and the peer-reviewed report is available at NIH NLM PUBMED PMID: 34546975 .

