## Supplemental Materials for "IL-6 trans-signaling regulates CXCL12 expression in mesenchymal stromal cells and plasma cell niche in rejecting lung allografts"

Vibha N. Lama, M.D., M.S.,

Division of Pulmonary, Allergy, Critical Care, and Sleep Medicine,

Department of Medicine Emory University School of Medicine;

101 Woodruff Circle, Atlanta, GA, USA 30322.

(404) 712-2790,

**One Sentence Summary:** We characterize CXCL12-expressing mesenchymal cells and their role in a pro-survival niche for antibody-secreting cells in rejecting lung allografts.

***RESULTS.***

**Table S1. Post-hoc histological analysis for the presence of plasma cells from normal and CLAD human explants.**

| **Slide** | **Diagnosis** | **Plasma Cells (-/+/++/+++)** | **Histological Findings** |
| --- | --- | --- | --- |
| S1 | RAS | **+ +** | Dense, hyalinizing pleuritis; fibroelastosis of proximal bronchovascular area; intimal fibrosis of large arteries; patchy chronic inflammation |
| S2 | RAS | **+ +** | Dense fibroelastosis of pleural and subpleural parenchyma; patchy chronic bronchovascular inflammation; focal obliterative bronchiolitis; intimal fibrosis of large arteries |
| S3 | RAS | **+ + +** | Prominent perivascular collagenous fibrosis, minimal chronic inflammation of peripheral lung |
| S4 | RAS | **+ + +** | Subpleural, pleural fibroelastosis; diffuse OB; focal atelectasis, fibroelastosis; mild chronic inflammation |
| S5 | RAS | **+ + +** | Perivascular and peribronchial fibrosis in large hilar area; dense collagenous pleuritis with chest wall adhesions; brisk chronic inflammation band in subpleura. |
| S6 | RAS | **+ + +** | Dense, fibrotic pleuritis with fibroelastosis; scattered chronic inflammatory aggregates (lymphocytes, plasma cells) predominantly adjacent to parenchymal border of pleural fibrosis; subepithelial fibrosis of bronchioles; intimal fibrosis of large arteries |
| S7 | RAS | **+ + +** | Fibrotic pleura with fibroelastosis and marked linear chronic inflammation layer (plasma cells and lymphocytes) adjacent to lung parenchyma; small to medium arterial and intimal fibrosis; foci of early acute lung injury |
| S8 | RAS | **+ + +** | Partial atelectasis; minimal chronic inflammation of peribronchial area; fibrous, organizing pleuritis with foci of fibroelastosis; linear band of chronic inflammation affecting subpleural area with plasma cells; intimal fibrosis in medium to large arteries |
| S9 | BOS | **-** | Acute and organizing DAD; organizing pneumonia; organizing acute fibrinous pleuritis |
| S10 | BOS | **+** | Diffuse pleuritis with PPFE-type morphology; early organizing fibrosis around bronchovascular area |
| S11 | BOS | **+** | Organizing diffuse alveolar damage with areas of early fibrosing DAD; focal areas of fibrosis suggestive of subpleural atelectasis |
| S12 | BOS | **+ +** | Acute DAD with organizing pneumonia; organizing pleuritis and chronic, fibrotic pleuritis; lymphoplasmacytic infiltrate in the border between organizing, fibrotic pleuritis |
| S13 | Normal | **-** | Congestion; acute bronchitis; patchy interstitial pneumonitis |
| S14 | Normal | **-** | Congestion; acute bronchitis; patchy interstitial pneumonitis; edema |

***Footnote:*** *CLAD, chronic lung allograft dysfunction; RAS, Restrictive allograft syndrome; BOS, Bronchiolitis obliterates syndrome; Normal, human lungs, no transplant; -, no plasma cells; +, few plasma cells; ++, many plasma cells; +++, profuse plasma cell infiltration; OB, obliterative bronchiolitis; DAD,*

*diffuse alveolar damage; PPFE, pleuroparenchymal fibroelastosis.*

***Fig. S1. Characterization of CXCL12^+^ MCs in Cxcl12^iCreERT2^Rosa26^tdTomato^ mice and in RAS allografts.*** *(****A****) Representative FACS gating scheme used to isolate cell CD45^–^CD31^–^EpCAM^–^PDGFRα^+^CXCL12^+/–^ MCs from Cxcl12^iCreERT2^Rosa26^tdTomato^ mice. (****B****) mRNA expression analyzed by real-time PCR of sorted populations from A. Values are represented as means ± SEM. *p<0.05, **p<0.01, ****p<0.0001. One-way ANOVA; post-hoc test: Bonferroni’s. (****C****) Immunofluorescent imaging of representative tissue section from Cxcl12^iCreERT2^Rosa26^tdTomato^ mice in the pleural region showing CXCL12^+^ MCs (red). (****D****) Immunofluorescent imaging of representative tissue section from day 28 murine RAS allografts in the pleural region showing infiltration of ASCs (CD138, red; IgG, green; co-localization, yellow) and B cells (B220, white) in the subpleural niche. n=3. Original magnification = 200x. Scale bars = 50 µm.*

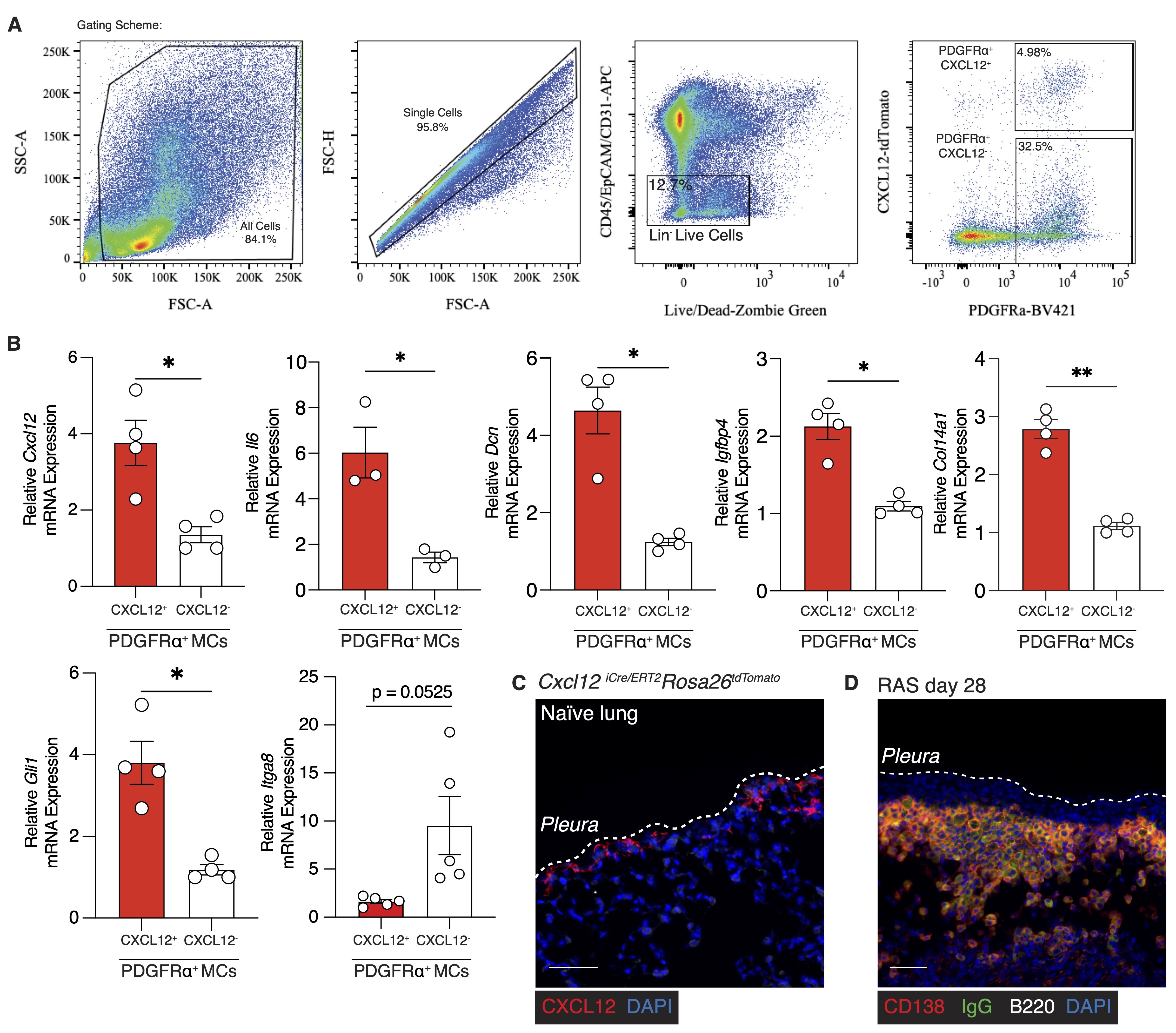

***MATERIALS AND METHODS.***

***Serum alloantibody assay.*** Using a modified protocol as previously described (*1*), 200 μL of PBS containing 0.5% BSA and 0.02% sodium azide was used to suspend 2 × 10^6^ thymocytes of donor origin (DBA/2J for isografts and RAS model, and C57BL/6J for BOS model) were mixed with 200 μL of serially diluted serum for 1 hour at 4°C with frequent agitation. After 3 washes with PBS, cells were stained for 30 minutes at 4°C with 100 μL of PBS containing 1 μL of polyclonal fluorochrome–conjugated goat anti–mouse IgM (μ chain specific) or anti–mouse IgG (Fcγ fragment specific) (Jackson ImmunoResearch; Cat#115-116-075,115-095-071, respectively). Cells were analyzed on a FACScan (BD Biosciences), and the median fluorescent intensity was calculated on FlowJo software (*ver.10.10.0*, Becton Dickinson, Covington, GA).

***Immunohistochemical staining of B cells in human and murine explants.*** Briefly, 5 µm FFPE sections were deparaffinized and rehydrated prior to immunostaining. H&E staining was performed using the Hematoxylin and Eosin Stain Kit (Cat#H-3502, Vector Laboratories, Newark, CA) according to the manufacturer’s instructions. Masson’s trichrome staining was performed using the Masson’s Trichrome Staining Kit (Cat#HT15-1KT, Sigma-Aldrich, St. Louis, MO). For immunofluorescence staining, sections were subjected to heat induced antigen retrieval (Citrate-based Antigen Unmasking Solution; Cat#H-3300-250, Vector Laboratories, Newark, CA) using a pressure cooker. Sections were then quenched for endogenous peroxide activity (0.3% hydrogen peroxide in methanol x 10 minutes), followed by protein block (2.5% goat serum in 0.5% PBS-Tween-20 x 1h). All primary antibodies were incubated overnight in a humid chamber at 4°C: anti-mouse B220 (1:100; Cat#ab233273, Abcam), anti-CD138 (1:250; Cat#LS-B8176-50, LS Biosciences), anti-mouse Ig-Cy3 and Ig-FITC (1:100; Cat#1010-01, Southern Biotech), anti-human Ig-HRP (1;100; Cat#2010-05, Southern Biotech), and anti-⍺SMA (1:200; Cat#2010-05, ThermoFisher Scientific). Appropriate secondary antibodies were incubated for one hour at room temperature using HRP-conjugated anti-mouse and anti-rabbit secondary antibodies (1:200; Cat#A8924 and 1:200; Cat#A0545, Sigma Aldrich, respectively), followed by Tyramide-Signal Amplification was utilized for unconjugated primary antibodies (NEL745001KT for Cy5, NEL741001KT for Fluorescein, and NEL744001KT for Cy3, Akoya Biosciences). Imaging was performed with a Nikon Eclipse Ti Fluorescent Microscope. Full slide scans were scanned on an Akoya Biosystems PhenoImager HT and analyzed using the QuPath software.

***Whole lung digestion.*** For mesenchymal cell isolation, lung explants were harvested, perfused, and digested via collagenase A at 37°C for 30 minutes in serum-free DMEM. Digests were further sheared using an 18-gauge needle and incubated for an additional 10 minutes at 37°C and then filtered through a 70 µm filter. Digestion reaction was stopped using DMEM containing 10% FBS followed by centrifugation (400xg for 10 minutes) and resuspension in FACS buffer.

***Isolation of mouse organs and single cell suspensions.*** Grafted lungs were perfused with PBS and harvested at days 7, 14, 28, and 40 post-transplant, minced and homogenized in 10 mL of FACS buffer using gentleMACS Dissociator (130-0930235, Miltenyi Biotech). The resulting suspension was passed through a 70 µm strainer, and cell counts were recorded prior to centrifugation (400xg for 10 minutes). 1x10^6^/mL concentration was used for FACS analyses.

For isolation of splenic and mesenteric lymph nodes cells, harvested organs were placed onto a 70 µm strainer and gently homogenized, and then washed using FACS buffer. Cell counts were recorded, and the strained cells were centrifuged at 400xg for 10 minutes.

***Flow cytometry analysis and cell sorting.*** For flow cytometry analysis, CD45, CD31, EpCAM, PDGFRα, CXCL12, CD19, CD38, CD3e, CD4, CD138, B220, Aqua Live/Dead, etc. (**Table S2**), were used for flow cytometry analysis. All gates are based off fluorescence minus one controls and single stains. Flow sorting was performed by either removing CD45, EpCAM, CD31-positive cells followed by gating for PDGFRα^+^ CXCL12-PE^+^ cells, or by gating all live cells and sorting CD45^+^, CD31^+^, EpCAM^+^, and PDGFRα^+^ cells. Sorted cells were sorted into tubes and their total RNA was immediately extracted using the QIAGEN RNeasy Mini kit according to the manufacturer’s protocol.

**Table S2. List of antibodies used in FACS analyses**

| **Target** | **Conjugate** | **Clone** | **Vendor** | **Cat#** | **Experiment** |
| --- | --- | --- | --- | --- | --- |
| CD138 | PE | W20051E | BioLegend | 112503 | FC |
| CD4 | BV711 | RM4-5 | BioLegend | 100557 | FC |
| CD8a | BV711 | 53-6.7 | BioLegend | 100759 | FC |
| Anti-Mouse Ig(H+L) | AF488 | — | Southern Biotech | 1010-30 | FC, IHC |
| CD19 | PE-Cy7 | ID3/CD19 | BioLegend | 152418 | FC |
| B220 | BV650 | RA3-6B2 | BioLegend | 103239 | FC |
| CD45 | APC | 30-F11 | BioLegend |  | FC |
| CD38 | APC-Cy7 | 90 | BioLegend | 102717 | FC |
| Ki-67 | BV421 | 16A8 | BioLegend | 652411 | FC |
| PDGFRα | BV421 | APA5 | BioLegend | 135923 | FC |
| EpCAM | APC | G8.8 | BioLegend | 118213 | FC |
| CD31 | APC | W18222B | BioLegend | 160209 | FC |
| Anti-Mouse IgG | BV421 | — | Jackson Immunoresearch | 115-675-071 | FC |
| Anti-Mouse IgM | APC | — | Jackson Immunoresearch | 115-136-075 | FC |
| Anti-RFP (Rb) | - | — | Rockland | 600-401-379 | IHC |
| CD138 | - | B‑A38 | LS Bio | LS‑B9360 | IHC |
| CD38 | HRP | H-11 | Santa Cruz | sc-374650 | IHC |
| Anti-Human IgG(H+L) | HRP | — | Southern Biotech | 2014-05 | IHC |
| B220 | - | RA3-6B2 | BioLegend | 103202 | IHC |
| GFP | FITC | — | Abcam | ab6662 | IHC |
| CCSP | - | CCSP his fusion | Seven Hills Bioreagents | WRAB-3950 | IHC |
| Alpha-SMA | - | 1A4 | Invitrogen | 14-9760-82 | IHC |
| E-Cadherin | - | ####### | Cell Signaling | 3195S | IHC |
| Phospho-Stat3 (Tyr705) | - | D3A7 | Cell Signaling | #9145 | WB |
| Stat3 | - | 124H6 | Cell Signaling | 9139 | WB |

Abbreviations: FC, Flow Cytometry; IHC, Immunohistochemistry; WB, Western Blot

***RNA isolation and Real-Time PCR.*** Total RNA was isolated using the RNeasy Mini Kit, (74104, QIAGEN) and cDNA was synthesized using the High-Capacity cDNA reverse transcription kit (4368814, Applied Biosystems). Real-time PCR was performed using human and mouse Taqman assays (**Table S3**), and TaqMan Gene Expression Master Mix (Cat#4369016, Applied Biosystems); and SYBR Green PCR Master Mix (4309155, Applied Biosystems) for mouse *Itga8* (FW: CCGAAGGCCAAGGTTACTG, RV: AACTTCCAGGTCCTCCCACT) with *Gapdh* as a control (FW: GTCAGCAATGCATCCTGCA, RV: CCGTTCAGCTCTGGGATGAC). The relative mRNA levels of the target genes were calculated as equal to 2^-(ΔCt target mRNA - ΔCt Β-Actin)^.

**Table S3. List of primers used in human and mouse real-time PCR analyses.**

| **Gene name** | **Mouse** | **Human** |
| --- | --- | --- |
| *Cxcl12/CXCL12* | Mm00445553_m1 | Hs00171022_m1 |
| *Il6/IL6* | Mm00446190_m1 | Hs00174131_m1 |
| *STAT3* | - | Hs00374280_m1 |
| *Actb/ACTB* | Mm02619580_m1 | Hs01060665_m1 |
| *Gli1* | Mm00494654_m1 | - |
| *Dcn* | Mm00514535_m1 | - |
| *Igfbp4* | Mm00494922_m1 | - |
| *Col14a1* | Mm00805269_m1 | - |
| *Inta8* | Mm01324958_m1 | - |

***Single-cell RNA sequencing analysis.*** Lungs from *Gli1^CreERT2/WT^*;*Rosa26^mTmG/WT^* B6D2F1/J mice were harvested. Single cell suspension was sorted for CD45^−^CD31^−^ cells by flow cytometry and analyzed by single-cell RNA-sequencing using the 10X Genomics Chromium Next GEM Single Cell 3′ Kit v3.1 (part number 1000268) following the manufacturer’s protocol. Bioinformatics analyses was previously reported (*2*), and re-examined for *Cxcl12 and Il6* gene expression.

***Primary mesenchymal cell isolation and culture conditions.*** Cellular fraction of the bronchoalveolar lavage fluid (BALF) utilized for MC isolation, were derived from lung transplant recipients diagnosed free of CLAD or having CLAD (either BOS or RAS). CLAD phenotype was identified based on ISHLT guidelines (*3*) and as previously described (*4, 5*). CLAD onset date was designated as the first date of spirometric decline, and CLAD was defined as BALF obtained ≤90 days before CLAD or after CLAD onset. Lung transplant recipients were designated CLAD-free in case of no evidence of acute rejection or infection. MCs were isolated and characterized as previously described (*6*). Briefly, cellular fraction from BALF was cultured until the formation of fibroblastoid colony-forming units (CFUs) in the growth media - Dulbecco’s Modified Eagle Medium (DMEM) with high glucose and glutamine (Invitrogen, Waltham, MA; Cat#11965) supplemented with penicillin/streptomycin (100 units/mL) and amphotericin B (0.5%). Cells at passage 3-6 were used for all experiments, wherein they were cultured to ~70% confluence and quiesced overnight prior to treatment. Cells were then treated in serum-free DMEM with human recombinant IL-6 (50 ng/mL; Cat#206-IL-200/CF), or human recombinant sIL-6R (200 ng/mL; Cat#227-SR-025/CF) from RnD Systems, Ruxolitinib (RUX; 100 nM), WHP154 (WHP; 100 nM), Actinomycin D (ActD; 5 μg/mL) from Selleckchem, Houston, TX, for 24 and 48 hours to collect RNA and whole cell lysates, respectively.

***CXCL12 ELISA.*** Supernatants from cell culture experiments treated for 48 hours in serum-free media were centrifuged at 10,000g for 10 minutes at 4°C to remove sediment and debris before use. CXCL12 levels were measured using the Human CXCL12/SDF-1⍺ Quantikine ELISA Kit (DSA00, R&D Systems, Minneapolis, MN) according to the manufacturer’s protocol. Absorbance at 450 nm was measured using a SpectraMax M3 multi-mode microplate reader (Molecular Devices, San Jose, CA).

***Immunoblot analysis.*** Whole cell lysates of MCs were extracted as previously reported (*7*). Protein concentrations were determined using a Pierce Coomassie Plus (Bradford) Assay Kit (23236, ThermoFisher Scientific) and a Thermo Scientific BioMate 3 Spectrophotometer. Western blotting was performed to analyze protein expression using primary antibodies directed against Phospho-STAT3 (Try705) (9145, Cell Signaling, 1:2000), and STAT3 (12640, Cell Signaling Technology, Danvers, MA, 1:1000). HRP-conjugated anti-rabbit secondary antibodies were A0545 (Sigma Aldrich, 1:10,000), respectively.

***Plasmid transfection.*** The control plasmid pFUGW was a gift from David Baltimore (Addgene plasmid #14883; RRID:Addgene_14883) and the EF.STAT3C.Ubc.GFP plasmid was a gift from Linzhao Cheng (Addgene plasmid #24983; RRID:Addgene_24983). Plasmids were amplified by transforming Stbl3 One Shot Chemically Competent E. coli (Cat#C737303, ThermoFisher Scientific) at 42°C in S.O.C. medium and plated on ampicillin-selective growth plates at 100 µg/mL. Single CFUs were selected and amplified by broth cultures overnight. Bacterial cultures were harvested using the QIAGEN Plasmid Midi Kit (Cat#12143) and isolated plasmid sequences were verified by standard sequencing methods. Purified plasmids were packaged into lentiviral vectors by the Emory Vector core, for use in mammalian cell culture. Briefly, cells were plated at 50% confluence in 6-well plates, followed by infection in serum-free medium with protamine sulfate as a linker. 24 hours post infection, fresh serum-containing medium was added for 48 hours before harvesting the cells for RNA and whole cell lysates.

***Lymphocyte migration assay.*** Conditioned media from IL-6 trans-signaling experiments was used as the chemoattractant to stimulate migration of lymphocytes through the 5.0 µm polycarbonate membrane contained in cell-culture inserts. 0.5 x 10^6^ splenocytes were suspended in 100 µL and placed in the 24-well cell culture insert. The lower chamber was filled with 500 µL of conditioned media from trans-signaling-treated MCs. Three hours-post exposure, the lymphocytes in the lower chamber were subjected to FACS analyses to determine the number of CD3^-^CD19^+^ B lymphocytes that had migrated through the membrane.

***Hydroxyproline Assay.*** Lung explants were homogenized in 1 mL of PBS containing protease inhibitor. 1 mL of 12N HCl is added to the homogenates followed by hydrolysis at 120°C for 24 hours. A total of 5 μL of each sample is combined with 5 μL citrate/acetate buffer (238 mmol/L citric acid, 1.2% glacial acetic acid, 532 mmol/L sodium acetate, 85 mmol/L sodium hydroxide) in a 96-well plate. A total of 100 μL of the solution chloramine T (0.282 g chloramine T to 16 mL of citrate/acetate buffer, 2.0 mL of n-Propanol, 2 mL double-distilled H2O) is then added for 30 minutes and incubated at room temperature. Then 100 μL of Ehrlich’s reagent (2.5 g paradimethylamino benzaldehyde added to 9.3 mL of n-Propanol, 3.9 mL of 70% perchloric acid), and incubated at 65°C for 30 minutes. The absorbance of each sample is measured at 550 nm using a SpectraMax M3 multi-mode microplate reader (Molecular Devices, San Jose, CA). Standard curves were generated using known concentrations of the hydroxyproline standard (Millipore Sigma, St. Louis, MO).
